# NEDD4 Regulates Ubiquitination and Stability of the Cell adhesion Molecule IGPR-1 via Lysosomal Pathway

**DOI:** 10.1101/2021.02.07.430113

**Authors:** Linzi Sun, Razie Amraei, Nader Rahimi

## Abstract

The cell adhesion molecule immunoglobulin and proline-rich receptor-1 (IGPR-1) regulates various critical cellular processes including, cell-cell adhesion, mechanosensing and autophagy. However, the molecular mechanisms governing IGPR-1 cell surface expression levels remains unknown. In the present study, we used an *in vitro* ubiquitination assay and identified ubiquitin E3 ligase NEDD4 and the ubiquitin conjugating enzyme UbcH6 involved in the ubiquitination of IGPR-1. *In vitro* GST-pulldown and *in vivo* co-immunoprecipitation assays demonstrated that NEDD4 binds to IGPR-1. Over-expression of wild-type NEDD4 downregulated IGPR-1 and deletion of WW domains (1-4) of NEDD4 revoked its effects on IGPR-1. Similarly, knockdown of NEDD4 increased IGPR-1 levels in A375 melanoma cells. Furthermore, deletion of 57 amino acids encompassing polyproline rich (PPR) motif on the C-terminus of IGPR-1 nullified the binding of NEDD4 with IGPR-1. Moreover, we demonstrate that NEDD4 promotes K48- and K63-dependent polyubiquitination of IGPR-1. The NEDD4-mediated polyubiquitination of IGPR-1 stimulated lysosomal degradation of IGPR-1 as the treatment of cells with the lysosomal inhibitors, bafilomycine and ammonium chloride increased IGPR-1 levels in the HEK-293 cells ectopically expressing IGPR-1 and in multiple human skin melanoma cell lines. Hence, these findings suggest that ubiquitin E3 ligase NEDD4 is a key regulator of IGPR-1 with a significant implication in the therapeutic targeting of IGPR-1.

## Introduction

The cell adhesion molecule immunoglobulin and proline-rich receptor-1 (IGPR-1, also known as TMIGD2/CD28H) is expressed in human endothelial and epithelial cells, including, skin melanocytes [1]. IGPR-1 modulates the key angiogenic responses in endothelial cells including capillary tube formation, and barrier function [2, 3]. IGPR-1 supports colon cancer cell growth in cell culture, and in mouse tumor xenograft [4]. More importantly, IGPR-1 senses and reacts to cellular environment by acting as a mechanosensing receptor [5]. For example, IGPR-1 responds to various cellular stresses and undergoes phosphorylation in response to flow shear stress [5] exposure to chemotherapeutic agent, doxorubicin [4, 6], and nutrient starvation [7]. Phosphorylation of IGPR-1 at Ser220 is mediated by Ser/Thr kinases, AKT [5] and IKKβ [8]. IKKβ-dependent activation of IGPR-1 mediates autophagy by a mechanism that involves activation of AMPK [7]. The extracellular domain of IGPR-1 contains a single Ig domain followed by a single transmembrane domain and intracellular domain with multiple poly-proline-rich (PPR) motifs [2, 9]. PPR motif is known to interact with the WW (tryptophan-tryptophan) and SH3 (Src homology-3) domains [10]. The interaction of IGPR-1 with the SH3 domain containing proteins was previously reported [1]. The recent studies revealed that the ligand recognition by WW domains are highly similar to that of SH3 domain [11, 12]. At the core of the WW and SH3 domains there is a highly conserved aromatic groove, which recognizes PPR motif [13]. The observed findings suggest a possible interaction between IGPR-1 and the WW domain-containing proteins. However, whether IGPR-1 interacts with the WW domain containing proteins remains unknown.

Neural precursor cell-expressed developmentally downregulated gene 4 (NEDD4, also called NEDD4-1) is a distinct ubiquitin E3 ligase that is ubiquitously expressed in various human organs and tissues including, skin, liver, thyroid, kidney[14] and blood vessels[15]. NEDD4 is consists of a N-terminus C2 domain, four WW domains, and a C-terminus HECT-type E3 ligase domain [16]. While the C2 domain is involved in calcium and lipid binding/membrane localization, WW domains of NEDD4 recognizes PPR motif[17]. NEDD4 can elicit both oncogenic [18–21] and tumor suppressor [22, 23] activities in human cancers. It is over-expressed in some cancer types including, prostate, bladder, and colon [18, 24] and exerts its oncogenic or tumor suppressor functions largely via ubiquitination of proteins with tumor suppressor or oncogenic activities such as PTEN, MDM2, AKT, Myc and Ras [25–29]. Mechanistically, the dichotomous function of NEDD4 in human cancers is centered on its indiscriminate ability to recognize the PPR motifs which are widespread motifs in proteins with diverse functions [25]. Here, we performed a series of *in vivo* and *in vitro* experiments to evaluate the importance of NEDD4 in IGPR-1 function and identified IGPR-1 as a novel substrate of NEDD4. NEDD4 binds to, and mediates polyubiquitination of IGPR-1 leading to its lysosomal-dependent degradation.

## Materials and Methods

### Plasmids, shRNAs and Antibodies

Construction and retroviral expression of IGPR-1 in HEK-293 cells was previously described [5, 30]. HA-NEDD4/pcDNA (Cat#11426), WW domain truncated NEDD4-Myc-pcMV5, GST-fusion NEDD4-WW1 (cat#104208), GST-fusion WW3 (cat#104210), and GST-fusion NEDD4-WW4 (cat#104211) all were purchased from Addgene. The development and validation of rabbit polyclonal anti-IGPR-1 antibody previously reported [2]. K48-linkage specific polyubiquitin (D9D5) Rabbit mAb (cat#8081), K63-linkage specific polyubiquitin (D7A11) Rabbit mAb (#5621), HA-Tag antibody (C29F4) Rabbit mAb (cat#3724) and Myc-Tag (71D10) Rabbit mAb (cat#2278) all were purchased from Cell Signaling Technology. Anti-ubiquitin (FK2) (cat #04-263) was purchased from Millipore-Sigma. HA-tagged wild-type ubiquitin and lysine mutant constructs were originally provided by Ted Dawson (Johns Hopkins University, Institute for Cell Engineering, Baltimore, MD) and Cam Paterson (University of North Carolina) [31]. Human NEDD4 shRNA (sc-41079-SH) which is pools of three specific 19-25 nucleotide sequences in length purchased from Santa Cruz Biotechnology.

### *In vitro* IGPR-1 ubiquitination assay

*In vitro* IGPR-1 ubiquitination assays were performed using ubiquitination kit (Enzo Life Sciences, BML-UW9920-0001) following the manufacturer’s instructions. In brief, the assay was performed in 50μl reaction volume with the following components as indicated: 2μg of each recombinant biotinylated ubiquitin, 2.5μM E2 (a mixture of E2 enzymes: UbcH1, UbcH2, UbcH3, UbcH5a, UbcH5b, UbcH5c, UbcH6, UbcH7, UbcH8, UbcH10, UbcH13/Mms2) or 100 ng of purified GST-IGPR-1 and 1 μg GST-NEDD4. Following the reaction, the samples were analyzed by Western blot via Streptavidin-HRP.

### IGPR-1 degradation assay

HEK-293 cells expressing IGPR-1 were grown in 60 centimeter plates overnight, and transfected with NEDD3-HA or WW domain mutant NEED4 (ΔWW-NEDD4-Myc) or empty vector, respectively. After 48 h, the cells were lysed, and the degradation of IGPR-1 was determined by Western blot analysis using anti-IGPR-1 antibody, and GAPDH as a loading control respectively.

### IGPR-1 downregulation assay

Downregulation of IGPR-1 was performed as described previously [31]. Briefly, cells were pretreated with cycloheximide (60μg) and chased for various time points as indicated in the figure legend. Cells were lysed, and whole-cell lysates were subjected to Western blot analysis using anti-IGPR-1 antibody.

### Cell transfection

HEK-293 cells were grown in DMEM 10% FBS media to 60-70% confluency. NEDD4 plasmids or desired plasmids were transfected into HEK-293 cells via PEI (polyethylenimine). After 48 hours, cells were lysed and subjected to immunoprecipitation or Western blotting as described in the figure legends.

### Retrovirus production

pMSCV.puro vector containing IGPR-1 or other cDNA of interest was transfected into 293-GPG cells, and viral supernatants were collected for five days as previously described [32].

### Immunofluorescence Microscopy

HEK-293 cells expressing IGPR-1 alone or to together with NEDD4 were seeded onto coverslips and grown overnight in 60-mm plates to 80-100% confluence. The cells were washed once with PBS and fixed with freshly prepared 4% paraformaldehyde for 15 min at room temperature. After washing three more times with PBS, the cells were permeabilized with 0.25% Triton X-100 in Western rinse for 10 min at room temperature and then washed three times with PBS. For staining of IGPR-1, the cells were blocked with (1:1) BSA in Western rinse for 1 h and washed once in PBS, followed with incubation with polyclonal anti-IGPR-1 antibody (1:1000) for 1 h and detected with rabbit FITC-conjugated secondary antibody. The coverslips were mounted in Vectashield mounting medium with DAPI onto glass microscope slides. The slides were analyzed under a fluorescence microscope.

### Statistical analyses

Experimental data were subjected to Student t-test or One-way analysis of variance analysis where appropriate with representative of at least three independent experiments. p<0.05 was considered significant or as indicated in the figure legends.

## Results

### NEDD4 Binds to and Promotes Ubiquitination of IGPR-1

The intracellular C-terminus of IGPR-1 distinctively holds multiple poly-proline rich (PPR) motifs along with several serine phosphorylation sites with a potential to interact with the WW domain containing NEDD4 ubiquitin E3 ligase family proteins (**Figure 1A**). Among all the nine NEDD4 family proteins, NEDD4 commonly targets cell surface proteins for ubiquitination with PPR motifs[17]. Hence we hypothesized that NEDD4 could interact with IGPR-1 leading to its ubiquitination. NEDD4 has four WW domains (**Figure 1B**). To test our hypothesis, we generated a panel of GST-fusion of recombinant proteins corresponding to WW1, WW3 and WW4 domains of NEDD4 (**Figure 1B, 1C**) and tested their abilities to interact with IGPR-1 in an *in vitro* GST-pulldown assay. The result showed that WW4 domain selectively interacted with IGPR-1 expressed in HEK-293 cells (**Figure 1D**). Expression of IGPR-1 in HEK-293 cells is shown (**Figure 1E**). To confirm our observation, we also tested the *in vivo* binding of IGPR-1 with the full-length NEDD4. To this end, we co-expressed NEDD4-HA with IGPR-1-Myc in HEK-293 cells followed by immunoprecipitation with IGPR-1 antibody and subsequent immunoblotting with anti-HA antibody for NEDD4. NEDD4 was co-immunoprecipitated with IGPR-1 (**Figure 1F**), indicating that NEDD4 binds with IGPR-1 *in vitro* and *in vivo*.

**Figure 1:**
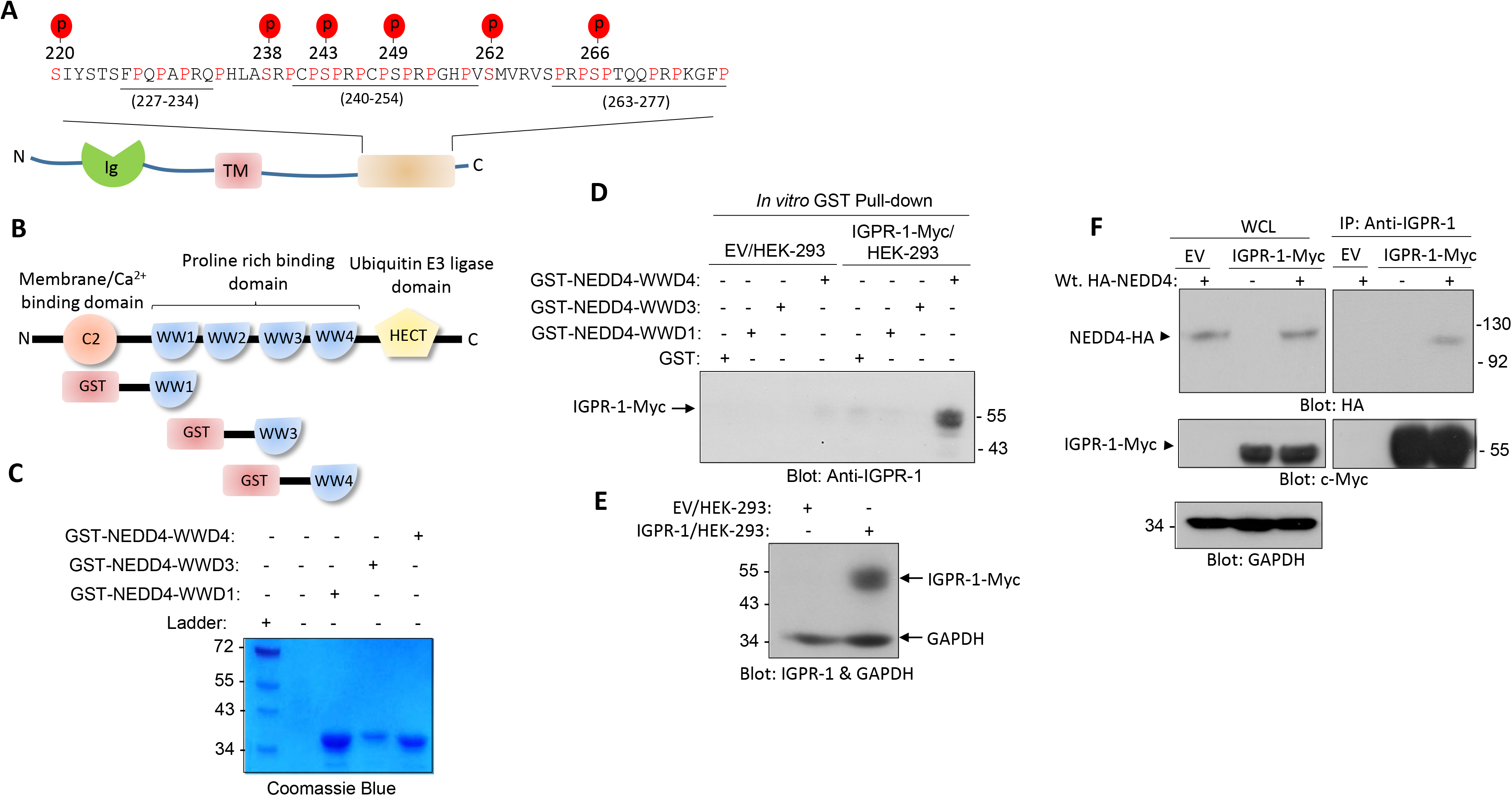
NEDD4 through its WW domain 4 binds to IGPR-1. (**A**) A schematic of IGPR-1 and the locations of the poly-proline rich motifs at the C-terminus. (**B**) Schematic of NEED4 domains and GST constructs corresponding to WW domains (#1,3,4). (**C**) Coomassie blue staining of purified GST-NEDD4 fusion WW domains. (**D**) HEK-293 cells expressing empty vector (EV) or IGPR-1-Myc were subjected to an *in vitro* GST-pull-down assay corresponding to WW domains (1, 3 and 4) of NEDD4, followed by Western blot analysis using an anti-IGPR-1 antibody. (**E**) Western blot analysis of whole cell lysate of HEK-293 cells expressing EV or IGPR-1-Myc immunoblotted for IGPR-1 or GAPDH. (**F**) HEK-293 cells expressing EV or IGPR-1-Myc were transfected with NEDD4-HA. After 48 hours post-transfection, cells were lysed and subjected to co-immunoprecipitation assay via anti-IGPR-1 antibody followed by Western blot analysis using HA antibody. The membrane was re-blotted with anti-IGPR-1 antibody. Whole cell lysate (WCL).

Next, we asked whether the presence of PPR motifs on IGPR-1 are responsible for the binding of IGPR-1 with NEDD4. To answer this question, we expressed the C-terminus truncated IGPR-1 (Δ57-IGPR-1) where the 57 amino acids compassing the key PPR motifs was deleted (**Figure 2A**). The whole cell lysates of HEK-293 cells expressing IGPR-1 and Δ57-IGPR-1 were subjected to an *in vitro* GST-pulldown assay, which showed that wild-type IGPR-1 but not Δ57-IGPR-1 binds to WW4-NEDD4 (**Figure 2B**). Expression of IGPR-1 and Δ57-IGPR-1 is shown **(Figure 2C**). Moreover, we examined the *in vivo* binding of NEDD4 with Δ57-IGPR-1 via a co-immunoprecipitation assay. Full-length NEDD4-HA interacted with wild-type IGPR-1 but not with Δ57-IGPR-1 (**Figure 2D**). Additionally, Western blot analysis of the whole cell lysates demonstrated that while over-expression of NEDD4 significantly reduced the full-length IGPR-1 levels, but it had no noticeable effect on Δ57-IGPR-1 levels (**Figure 2E**). Taken together, the data demonstrate that IGPR-1 via its PPR motifs interacts with the WW domain #4 of NEDD4 and that deletion of PPR motifs renders IGPR-1 insensitive to NEDD4.

**Figure 2.**
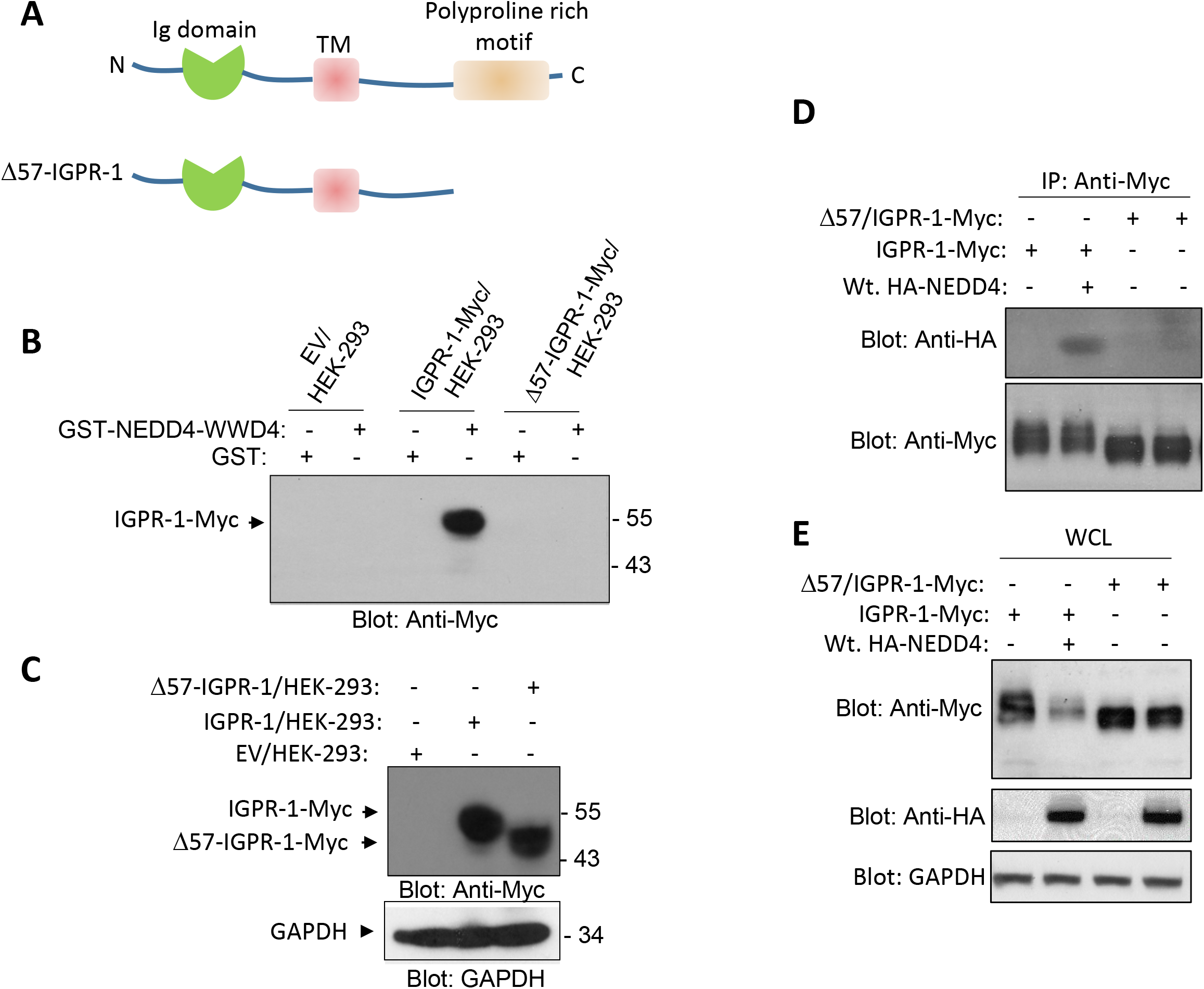
Deletion of proline-rich motifs on IGPR-1 eliminates its binding with NEDD4: (**A**) Whole cell lysates from HEK-293 cells expressing empty vector, IGPR-1-Myc or Δ57/IGPR-1-Myc were subjected to in vitro GST-pull down assay using GST-NEDD4-WW#4 followed by Western blot analysis using anti-Myc antibody. (**B**) Whole cell lysates corresponding to IGPR-1-Myc or Δ57/IGPR-1-Myc expressed in HEK-293 cells and loading control, GAPDH. (**C**) Whole cell lysates from HEK-293 cells expressing IGPR-1-Myc or Δ57/IGPR-1-Myc were subjected to co-immunoprecipitation assay using anti-Myc followed by Western blot analysis using anti-HA antibody. (**D**) Whole cell lysates corresponding to IGPR-1-Myc, Δ57/IGPR-1-Myc, HA-NEDD4 expressed in HEK-293 cells and loading control, GAPDH.

### NEDD4 mediates ubiquitination of IGPR-1

To test whether NEDD4 interaction with IGPR-1 mediates ubiquitination of IGPR-1, we used an *in vitro* ubiquitination assay consisting of NEDD4 as a source of ubiquitin E3 ligase, eleven E2 conjugating enzymes, biotinylated ubiquitin and GST-IGPR-1 encompassing the cytoplasmic domain of IGPR-1 as a substrate. The schematic of the *in vitro* ubiquitination assay is shown (**Figure 3A**). The result demonstrated that NEDD4 strongly catalyzes ubiquitination of IGPR-1 in the presence of E2 conjugating enzyme, UbcH6 (**Figure 3B**). NEDD4 also ubiquitinated IGPR-1 in the presence of E2 conjugating enzymes, UbcH5c, UbcH5b and UbcH5a, albite ubiquitination of IGPR-1 was significantly weaker compared to UbcH6 (**Figure 3B**). Expression of GST-NEDD4, GST-IGPR-1 and E2 conjugating enzymes are shown (**Figure 3C-E**). To investigate whether NEDD4 also ubiqutinates IGPR-1 *in vivo,* we expressed HA-NEDD4 in IGPR-1/HEK-293 cells and analyzed ubiquitination of IGPR-1 through immunoprecipitation followed by Western blotting assay using anti-ubiquitin (FK2) antibody, which detects both mono- and poly-ubiquitinated proteins. Expression of NEDD4 in IGPR-1/HEK-293 cells promoted ubiquitination of IGPR-1 as detected with anti-ubiquitin (FK2) antibody (**Figure 3F**). Further analysis demonstrated that NEDD4 stimulated both K63- and K48-dependent polyubiquitination of IGPR-1 as detected by K63- and K48-specific polyubiquitin antibodies (**Figure 3F**). Next, we asked whether Δ57-IGPR-1, which cannot bind to NEDD4 (**Figure 2B**), escapes from the NEDD4-medaited ubiquitination of IGPR-1. The result showed that while, wild-type IGPR-1 is strongly ubiquitinated by NEDD4, no significant ubiquitination was observed for Δ57-IGPR-1 (data not shown).

**Figure 3:**
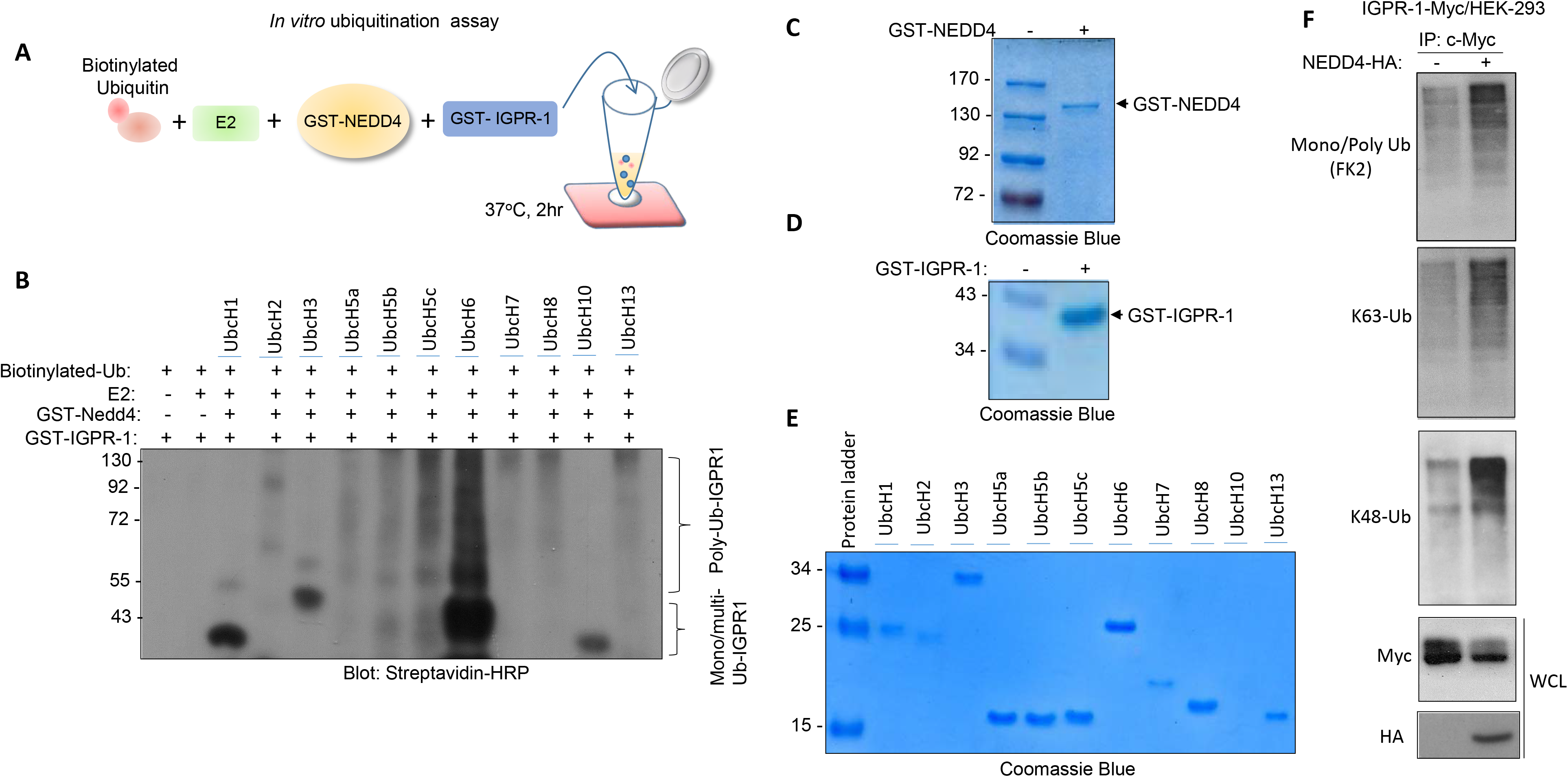
NEDD4 mediates mono- and polyubiquitination of IGPR-1. (**A**) A schematic of *in vitro* ubiquitination assay is shown. (**B**) *In vitro* ubiquitination assay was carried out as described in the material and method section using biotinylated ubiquitin (Ub), E2 conjugating enzymes, purified GST-NEDD4 and GST-cytoplasmic domain of IGPR-1. The samples were resolved on SDS-PAGE and ubiquitinated GST-IGPR-1 was detected by Western blotting via Streptavidin-HRP antibody. (**C-E**) Coomassie blue staining of GST-IGPR-1, GST-NEDD4 and E2 conjugating enzymes. (**F**) HEK-293 cells expressing IGPR-1-Myc were transfected with an empty vector or wild-type NEDD4-HA, cells were lysed and whole cell lysates were immunoprecipitated with an anti-Myc antibody followed by Western blot analysis using anti-ubiquitin (FK2) antibody, K63-poly-ubiqutin antibody or K48-poly-ubiqutin antibody. Whole cell lysates (WCL) were blotted for IGPR-1-Myc and NEDD4-HA. The graphs are representative of three independent experiments. P<0.05.

To probe further the role of specific lysine residues involved in the regulation of IGPR-1, we expressed HA tagged wild-type ubiquitin, KO mutant, lysine 48 (K48, ubiquitin with only a K48 residue, other lysine residues were mutated to arginine) or lysine 63 (K63, other lysine residues were mutated to arginine) ubiquitin constructs (**Figure 4A**) in IGPR-1/HEK293 cells and examined their effects on the downregulation of IGPR-1 via immunoprecipitation with anti-Myc (for IGPR-1) followed by Western blotting with anti-HA antibody (for ubiquitin). The result showed that HA tagged wild-type ubiquitin, lysine 48 (K48) and lysine 63 (K63) ubiquitin all were conjugated to IGPR-1 relatively in a similar manner (**Figure 4B**), indicating that K48- and K63-dependent ubiquitination could regulate degradation and downregulation of IGPR-1, respectively. Consistent with the known role of K48 in protein degradation[33], co-expression of K48-ubiqutin, as well as wild-type ubiquitin, but not K63-ubiqutin, with IGPR-1 significantly downregulated IGPR-1 levels (**Figure 4C**).

**Figure 4.**
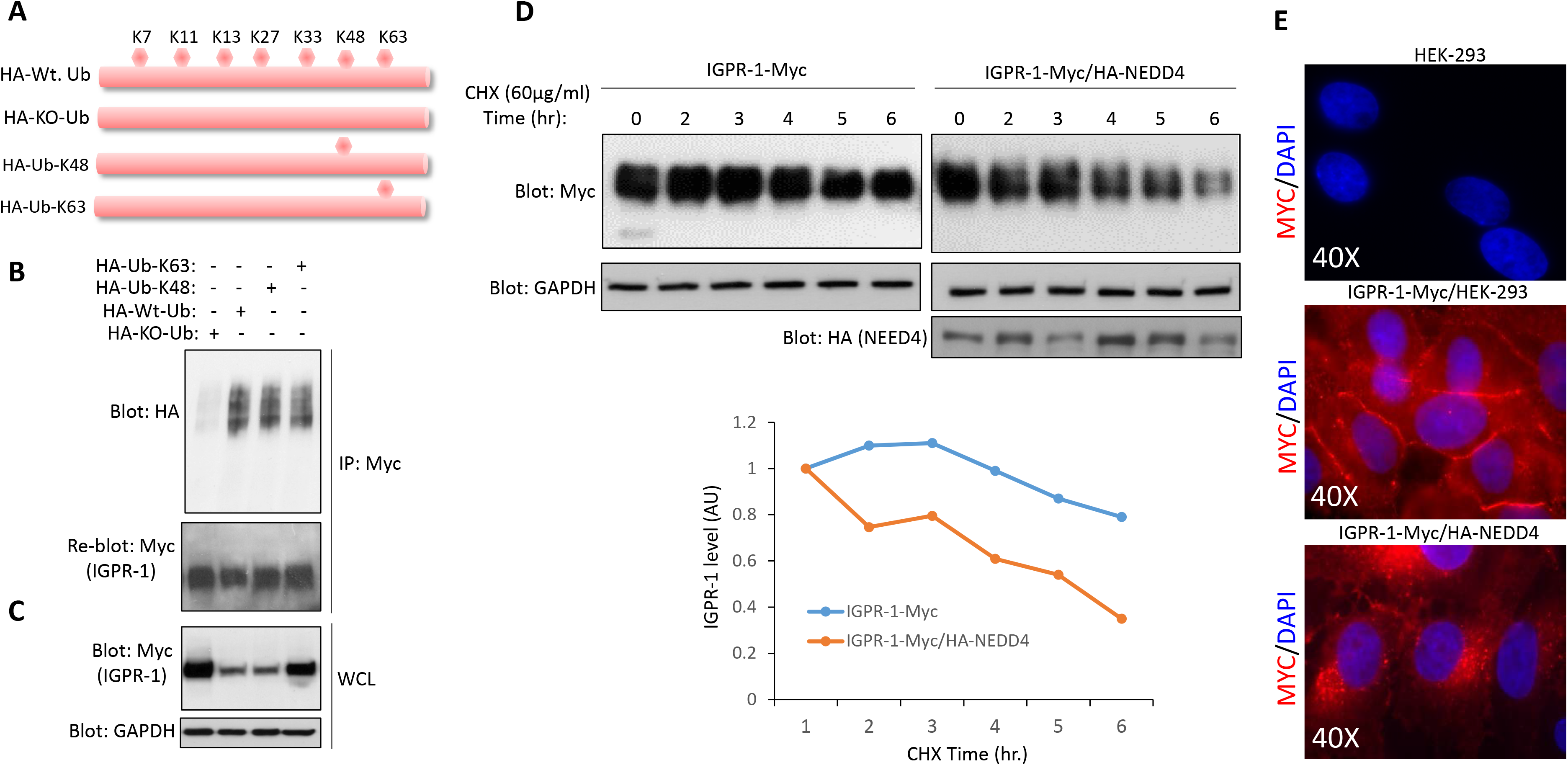
NEED4 mediates K48- and K63-dependent polyubiquitination and downregulation of IGPR-1. (**A**) A schematic of ubiquitin (Ub) constructs are shown. (**B**) HEK-293 cells expressing IGPR-1 were transfected with wild-type Ub, KO-Ub, K48-Ub or K63-Ub constructs. After 48 hours, cells were lysed and whole cell lysates were immunoprecipitated with anti-Myc and blotted with anti-HA antibody for ubiquitin. The same membrane was re-blotted for IGPR-1. (**C**) Whole cell lysates (WCL) was blotted for total IGPR-1, and loading control GAPDH. (**D**) IGPR-1/HEK-293 and IGPR-1/HEK-293 cells transfected with NEDD4 were subjected to cycloheximide pulse-chase assay. Cells were lysed at the indicated time points and whole cell lysates was subjected to Western blot analysis using anti-Myc antibody. Expression of NEDD4 and loading control GAPDH are shown. (**E**) IGPR-1/HEK-293 and IGPR-1/HEK-293 cells transfected with NEDD4 were stained with anti-Myc and subjected to immunofluorescence microscopy. 40X magnification.

### NEDD4 promotes lysosomal-dependent degradation of IGPR-1

Ubiquitination of cell surface proteins can promote downregulation (*i.e*., removal of receptor from cell surface via endocytosis) and degradation [33]. Therefore, we asked whether NEDD4-mediated ubiquitination of IGPR-1 promotes downregulation of IGPR-1. To answer this question, we first subjected IGPR-1/HEK-293 and IGPR-1/HEK-293 cells co-expressing NEDD4 to downregulation assay via cycloheximide pulse-chase analysis. Over-expression of NEDD4 significantly reduced the half-life of IGPR-1 (about 65% versus 11% at 6 hours chase) (**Figure 4D**). Furthermore, immunofluorescence staining analysis revealed that co-expression of NEDD4 with IGPR-1 greatly reduced the cell surface expression of IGPR-1 (**Figure 4E**), indicating that NEDD4 promotes downregulation of IGPR-1.

Next, we asked whether interfering with the interaction of NEDD4 with IGPR-1 inhibits its activity toward IGPR-1. To this end, we expressed wild type NEDD4 or ΔWW-NEDD4, where all the four WW domains of NEDD4 were deleted, in IGPR-1/HEK-293 cells and measured IGPR-1 levels via Western blot analysis. While over-expression of ΔWW-NEDD4 had no effect on the downregulation of IGPR-1, over-expression of wild-type NEDD4 markedly downregulated IGPR-1 (**Figure 5A**). Of note, in addition to NEDD4, over-expression of ITCH, a NEDD4 related HECT-type ubiquitin E3 ligase [34], also downregulated IGPR-1 (data not shown).

**Figure 5:**
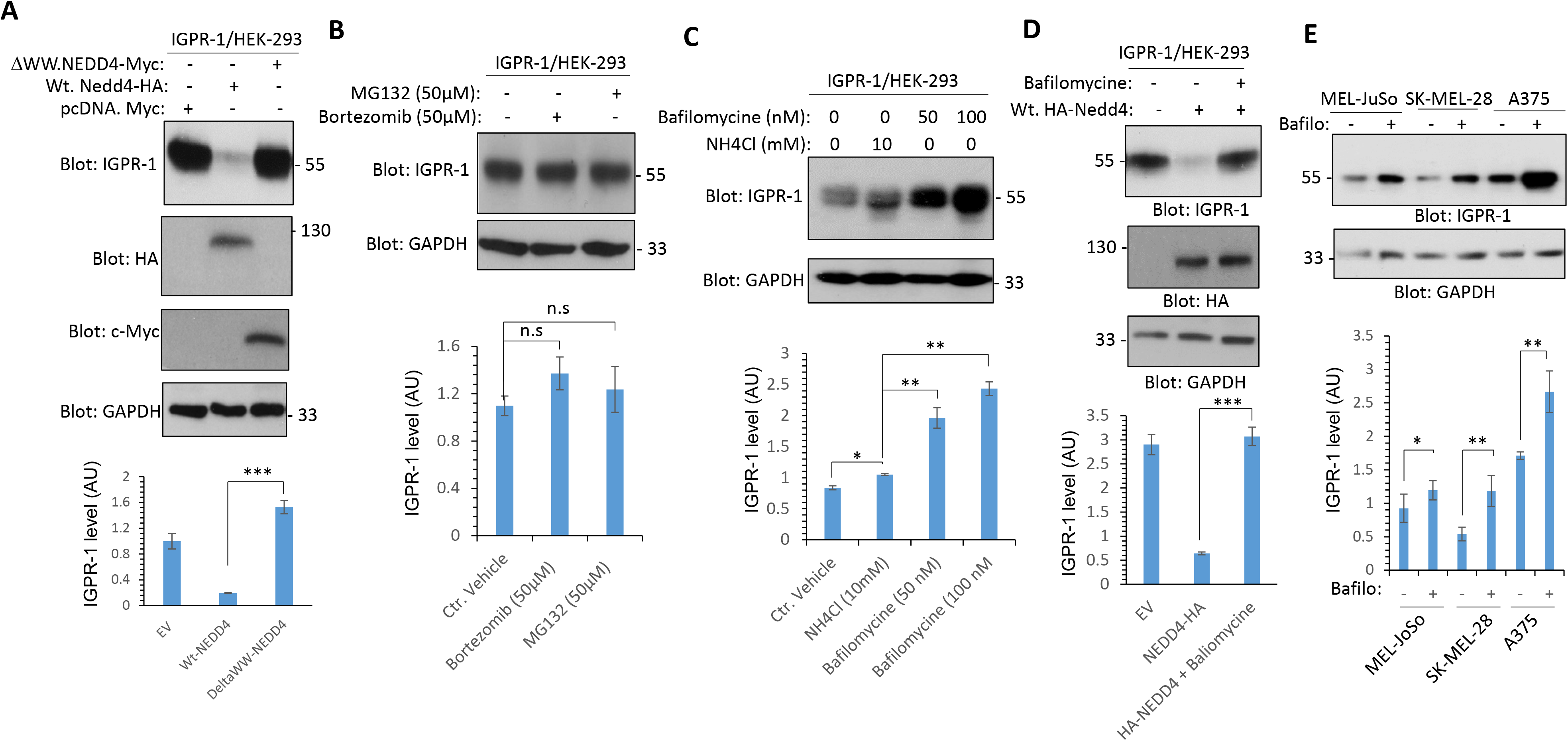
NEDD4 promotes lysosomal degradation of IGPR-1. (**A**) HEK-293 cells expressing IGPR-1 were transfected with an empty vector or wild-type NEDD4-HA or WW domain truncated NEDD4, where all the four WW domains was deleted (ΔWW-NEDD4). Cells were lysed and whole cell lysates were immunoblotted with an anti-IGPR-1 antibody, anti-HA antibody or Anti-Myc antibody for ΔWW-NEDD4. The membrane was also blotted for GAPDH for protein loading control. (**B**) HEK-293 cells expressing IGPR-1 were treated with MG132 or bortezomib for two hours, cells were lysed and whole cell lysates were blotted for IGPR-1 or GAPDH. (**C**) HEK-293 cells expressing IGPR-1 were treated with control vehicle or bafilomycine with different concentrations for 18 hours. Cells were lysed and whole cell lysates were blotted for IGPR-1 or GAPDH. (**D**) HEK-293 cells expressing IGPR-1 were transfected with an empty vector or wild-type NEDD4-HA. After 46 hours of transfection, cells were with control vehicle or bafilomycine and incubated for additional 18hours. Cells were lysed and whole cell lysates were blotted for IGPR-1 or GAPDH. (**E**) Melanoma cell lines (MEL-JoSo, SK-Mel-28 and A375) were treated with control vehicle or bafilomycine for 18 hours. Cells were lysed and whole cell lysates were blotted for IGPR-1 or GAPDH. The graphs (**A-E**) are representative of three independent experiments. P<0.05.

To further map the role of ubiquitination on IGPR-1 by NEDD4, we asked whether NEDD4-mediated downregulation of IGPR-1 is established through a mechanism that involves proteasomal-or lysosomal-dependent degradation. To test the involvement of proteasomal-pathway in the degradation of IGPR-1, we treated IGPR-1/HEK-293 cells with bortezomib or MG132 which both are cell-permeable proteasome inhibitors. Bortezomib and MG132 treatment had no major noticeable effects on IGPR-1 levels (**Figure 5B**), indicating that the 26S-proteasome pathway likely is not a major mechanism associated with the degradation of IGPR-1.

Because, inhibition of proteasome pathway did not alter IGPR-1 levels, we reasoned that lysosome-dependent pathway could likely control IGPR-1 levels. To this end, we treated IGPR-1/HEK-293 cells with NH4Cl or bafilomycine which are most commonly used lysosomal inhibitors. Treatment of cells with bafilomycine significantly increased IGPR-1 levels in a dose-dependent manner (**Figure 5C**), whereas NH4Cl (10mM) had only a minor effect on IGPR-1 levels (**Figure 5C**). However, at the higher concentration (20mM), NH4Cl increased IGPR-1 levels (data not shown). Next, we asked whether bafilomycine could blunt the effect of NEDD4 on IGPR-1. Over-expression of wild type NEDD4 in IGPR-1/HEK-293 cells downregulated IGPR-1 and treatment of cells with bafilomycine inhibited the NEDD4-dependent downregulation of IGPR-1 (**Figure 5D**), indicating that NEDD4 promotes lysosomal-dependent degradation of IGPR-1 in HEK-293 cells.

Considering the observed effect of bafilomycine on the ectopically expressed IGPR-1 in HEK-293 cells, we asked whether treatment of skin melanoma cells with bafilomycine could increase IGPR-1 expression endogenously in these cells. IGPR-1 is expressed in human skin melanocytes [2] and melanoma (our unpublished data). Treatment of MEL-JoSo, SK-MEL-28 and A375 melanoma cells with bafilomycine significantly increased IGPR-1 levels. The effect of bafilomycine on the IGPR-1 expression, particularly in A375 cells was more pronounced than the two other cell lines (**Figure 5E**). Taken together, the data demonstrate that NEDD4-mediated polyubiquitination on IGPR-1 leads to its lysosomal-dependent degradation.

### NEDD4 regulates expression of IGPR-1 in human skin melanoma

Considering that IGPR-1 is expressed at variable levels in human skin melanoma cell lines (**Figure 5E**), we decided to explore its expression profile in human skin melanoma. Our analysis of human skin melanoma dataset via cBioportal (http://www.cbioportal.org/) revealed that IGPR-1 mRNA levels is altered in 10.84% of patients (48 out of 443 cases) (**Figure 6A, B**). Interestingly, in 5.8% cases, IGPR-1 mRNA levels was elevated, whereas in 4.97% cases it was decreased (**Figure 6B, C**), suggesting a unique mechanism of alteration in the transcriptional regulation of IGPR-1 in the subset of human skin melanoma. Next, we asked whether there is a link between IGPR-1 protein levels and NEDD4 in human skin melanoma cell lines. Western blot analysis revealed that overall there is an inverse correlation between IGPR-1 and NEDD4 in four different melanoma cell lines (**Figure 6D**). Of note, IGPR-1 is not present in rodents [2] and hence B16F melanoma cells were negative for IGPR-1 and were used here as a negative control (**Figure 6D**).

**Figure 6.**
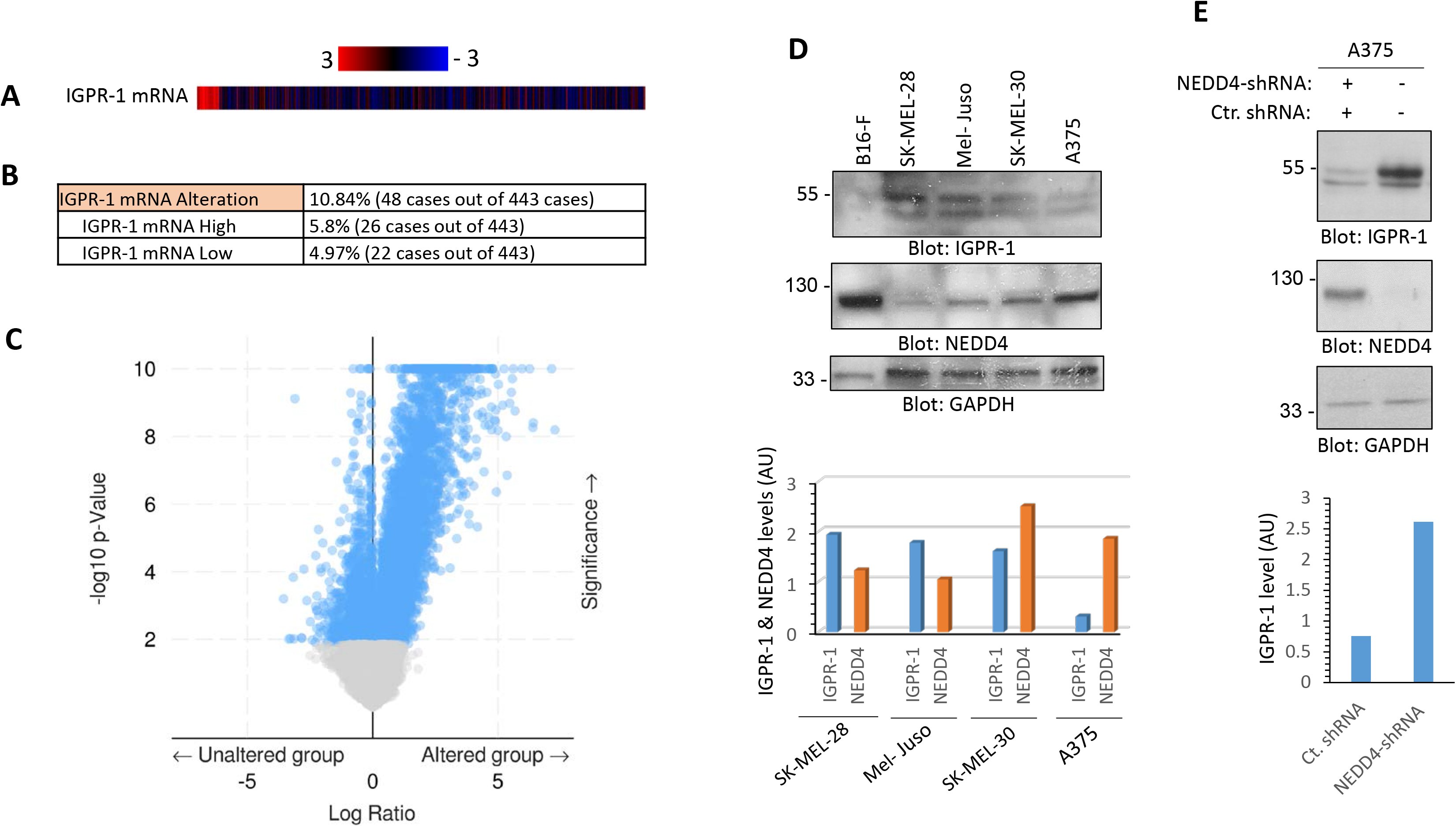
IGPR-1 is expressed in human melanoma and knockdown of NEDD4 increases IGPR-1 expression. (**A, B**) Expression of IGPR-1 in human melanoma. Data extracted from TCGA dataset via cBioportal (http://www.cbioportal.org/). (**C**) The volcano plot of IGPR-1 is shown. (**D**) Expression of IGPR-1 and NEDD4 in human melanoma cell lines. (**E**) A375 melanoma cells were transfected with human NEDD4 shRNA and after selection with puromycin cells were lysed and subjected Western blot analysis using anti-NEDD4, anti-IGPR-1 antibodies.

Next, we asked whether high levels of NEDD4 is a contributing factor for the relatively low levels of IGPR-1 in skin melanoma cells. To answer this question, we knockdown NEDD4 via shRNA strategy in A375 cell line which has highest levels of NEDD4 expression compared to three other melanoma cell lines (**Figure 6D**). NEDD4-shRNA almost completely knocked down NEDD4 (**Figure 6E**) and also significantly increased expression of IGPR-1 levels in A375 cells (**Figure 6E)**. Furthermore, re-expression of NEDD4 in these cells abrogated the effect of NEDD4-shRNA (data not shown). Taken together, the data demonstrate that NEDD4 plays a critical role in the regulation of IGPR-1 in human skin melanoma.

## Discussion

Ubiquitin modification can regulate endocytosis, recycling, and degradation of cell surface receptors including, ligand-stimulated receptors such as GPCRs and RTKs as well as cell adhesion molecules [33, 35–37]. In the present study, we have elucidated several key molecular features of IGPR-1 regulated by ubiquitination. We found a complex formation between WW domain 4 on NEDD4 and PPR motif on IGPR-1 drives ubiquitination of IGPR-1. NEDD4-mediated IGPR-1 polyubiquitination regulates receptor stability through lysosomal degradation pathway. IGPR-1 is expressed in the endothelial cells of vein and arteries, stimulates angiogenesis, regulates endothelial barrier function [2, 30, 38], and response to shear stress [5]. Moreover, IGPR-1 is expressed in human colon cancer and promotes tumor growth [4], skin melanoma cell lines (data shown in this manuscript) and induces autophagy with a significant implication in tumorigenesis and angiogenesis [8]. Identification of NEDD4 as an E3 ubiquitin ligase involved in the regulation of availability IGPR-1 at the cell surface represents a key mechanism with major implications in cell-cell adhesion, mechanosensing and tumor promoting activity of IGPR-1. Interestingly, NEDD4 promotes both K48- and K68-dependent polyubiquitination of IGPR-1 promoting both endocytosis and degradation of IGPR-1. Similar observations for NEDD4-mediated polyubiquitination of other proteins were previously reported [20, 39, 40].

Curiously, IGPR-1 expression in human skin melanoma is altered both at the mRNA and protein levels underscoring its potential role in tumorigenesis, which warrants further investigation. Similarly, NEDD4 expression and activity in various tumors including, skin melanoma is linked to tumorigenesis either by acting as a tumor suppressor or oncogene [25–29, 41]. NEDD4 is known to regulate cell proliferation, cell migration, and tumorigenesis in various cancer types [25]. The same cellular processes are also regulated by IGPR-1 [2, 4, 5, 8, 30]. In conclusion, NEDD4 together with ubiquitin conjugating enzyme, UbcH6 plays pivotal roles in regulating IGPR-1 ubiquitination, downregulation, and stability. These findings offer novel mechanistic insights for regulation of stability cell surface IGPR-1 and possible drug target for IGPR-1 associated pathologies such as angiogenesis and human cancers.

